# *Escherichia coli* can survive stress by noisy growth modulation

**DOI:** 10.1101/265801

**Authors:** Om Patange, Christian Schwall, Matt Jones, Douglas Griffith, Andrew Phillips, James C.W. Locke

## Abstract

Gene expression can be noisy^1,2^, as can the growth of single cells^3,4^. Such cell-to-cell variation has been implicated in survival strategies for bacterial populations^5–7^. However, it remains unclear how single cells couple gene expression with growth to implement these survival strategies. Here we show how noisy expression of a key stress response regulator, *rpoS*^8^, allows E. coli to modulate its growth dynamics to survive future adverse environments. First, we demonstrate that *rpoS* has a long-tailed distribution of expression in an unstressed population of cells. We next reveal how a dynamic positive feedback loop between *rpoS* and growth rate produces multi-generation *rpoS* pulses, which are responsible for the *rpoS* heterogeneity. We do so experimentally with single-cell, time-lapse microscopy^9^ and microfluidics^10^ and theoretically with a stochastic model^11,22^. Finally, we demonstrate the function of the coupling of heterogeneous *rpoS* activity and growth. It enables *E. coli* to survive oxidative attack by causing prolonged periods of slow growth. This dynamic phenotype is captured by the *rpoS*-growth feedback model. Our synthesis of noisy gene expression, growth, and survival paves the way for further exploration of functional phenotypic variability.

---

*E. coli* respond to stress by expressing a host of protective genes. Global stress response is controlled, in large part, by *rpoS,* which is an alternative sigma factor^8^. Sigma factors are a component of the RNA polymerase holoenzyme that recognise and bind to the promoter region of genes^13^. The housekeeping sigma factor, *σ*^70^, promotes the transcription of genes responsible for growth, for instance ribosomal genes^14^. On the other hand, *rpoS* up-regulates stress response genes^8^ (Fig. 1a). It is strongly up-regulated in the transition from exponential to stationary phase when cells are starved for resources^15^. Populations in exponential phase have been shown to express small amounts of functional *rpoS*^16,17^. However, these studies were of bulk cultures, which can mask single cell phenotypes.

**Figure 1.**
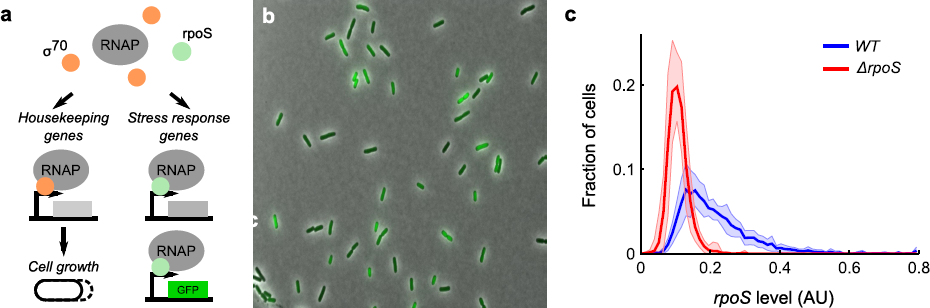
The stress response master regulator, rpoS, is heterogeneously expressed in unstressed cells. **a,** Schematic of the role of sigma factors δ^*70*^ and *rpoS* in promoting growth and activation of the stress response regulon, respectively. Also illustrated is the *rpoS* reporter, a transcriptional fusion to a stress response promoter. **b,** Representative phase and fluorescence composite image of *rpoS* reporter, *P_bo_i_A_-GFP,* in *WJ,* channel ranges chosen for display. **c,** Histograms of mean *rpoS* per cell (line: mean, shaded region: ± std dev) in *WJ* (10 biological replicates, 4,037 cells, mean = 0.21, CV = 0.51) and *ΔrpoS* (9 bio. reps., 4,069 cells, mean = 0.11, CV = 0.27) strains. The long tail of high *rpoS* levels present in the *WT* is absent in the knockout.

We therefore first asked the question: How is this small *rpoS* expression in exponential phase distributed amongst single cells? It could be that all cells have basal levels of *rpoS* or some cells could express the majority of the *rpoS.* To answer this question we grew cells in bulk culture into exponential phase and examined aliquots of the culture with single cell resolution under a microscope^9^ (see Fig. 1b, and Methods). As a proxy for *rpoS* we used a transcriptional reporter with a promoter from an rpoS-responsive gene fused to GFP: *P_bolA_-GFP^14^’^18^.* By computing histograms of mean *rpoS* level per cell we discovered that *rpoS* is heterogeneously distributed amongst single cells (Fig. 1c). To test our conclusion we carried out the same liquid culture assay on an rpoS-knockout *(ΔrpoS,* Fig. 1c). The characteristic long tail of the heterogeneous WT distribution vanished in the knockout strain, with gene expression levels near background. We found similar behaviour when alternative reporters for *rpoS* were tested (Sup. Fig. 1)^14^. To test whether the long-tail was specific to *rpoS,* we examined *σ*^70^ reporters. The distributions of *σ*^70^ levels in *WT* populations had less pronounced long-tails due to the higher abundance of *σ*^70^ in cells and did not change significantly in *ΔrpoS* (Sup. Fig. 2)^14^.

**Supplementary Figure 1.**
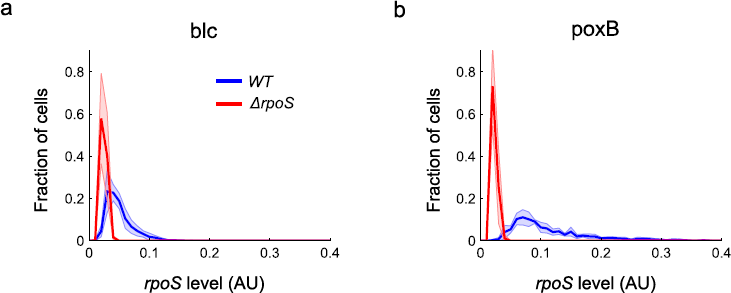
Alternative rpoS reporters have long-tailed distributions of rpoS levels; the long tails vanish in the rpoS-knockout. **a,** Transcriptional fusion of *P_bIc_-GFP* in *WT* (6 biological replicates, 2,509 cells, mean = 0.050 AU, CV = 0.46) and *ΔrpoS* (4 bio. reps., 1,190 cells, mean = 0.025 AU, CV = 0.21). **b,** Similarly for *P_poxB_-GFP (WT:* 5 bio. reps., 1,087 cells, mean = 0.12 AU, CV = 0.59; *ΔrpoS:* 7 bio. reps., 1,463 cells, mean = 0.023 AU, CV = 0.17). Lines and shaded region are mean ± std dev, respectively.

**Figure 2.**
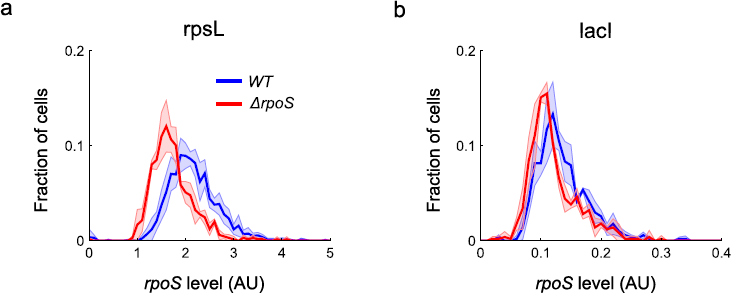
Reporters of *δ*^70^ have distributions with lower coefficients of variation than rpoS reporters and distributions that are similar in WT and ΔrpoS. **a,** Transcriptional fusion of *PrpsL-GFP* in WT (5 bio. reps., 1,576 cells, mean = 2.1 AU, CV = 0.25) and *ΔrpoS* (3 bio. reps., 647 cells, mean = 1.7 AU, CV = 0.25). **b,** Similarly for *P_tad_-GFP* in WT (3 bio. reps., 503 cells, mean = 0.14 AU, CV = 0.31) and *ΔrpoS* (3 bio. reps., 497 cells, mean = 0.12 AU, CV = 0.34). Lines and shaded region are mean ± std dev, respectively.

We next investigated the mechanism by which the *rpoS* distribution is produced. Reasoning that the distribution is due to a dynamic equilibrium, not a fixed subpopulation, we tracked single cells over multiple generations using time-lapse microscopy^9^ and the Mother Machine microfluidic device^10^ (Fig. 2a, Methods). Indeed, we found rich dynamic *rpoS* activity (see Methods). Some cell lineages have high *rpoS* activity pulses lasting multiple generations while others have very small pulses (Fig. 2b, Methods). We found a long-tailed distribution of pulse heights supporting the idea that the long-tailed liquid culture distribution is generated by cells pulsing *rpoS* on to different levels (Fig. 2c). We chromosomally integrated the *P_bolA_-GFP* reporter and found a similar consistency between bulk culture and microfluidic experiments suggesting the dynamics did not arise due to plasmid segregation noise (Sup. Fig. 3a-c). However, the fluorescence signal was very dim, thus we proceeded with the plasmid-based reporter.

**Figure 2.**
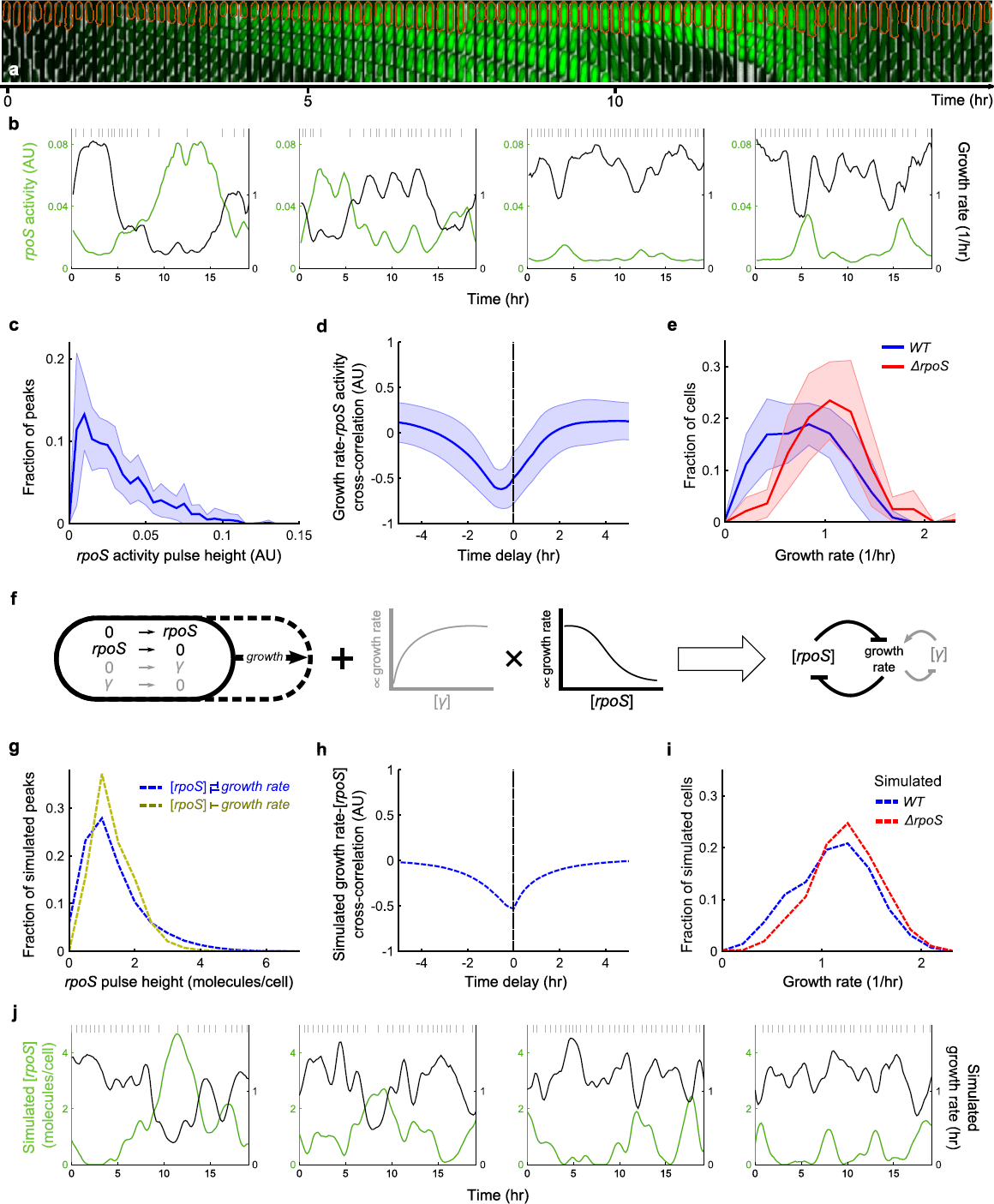
Growth-rpoS mutual inhibition produces multi-generation rpoS pulses and heterogeneous rpoS expression. **a,** Sample montage of a mother cell in the Mother Machine (orange outline) pulsing on *rpoS* and reducing growth rate. Phase contrast and fluorescence channel ranges chosen for display. **b,** Sample time traces of *rpoS* activity and growth rate for four mother cells. Grey vertical lines indicate cell divisions. **c**, Histogram of *rpoS* activity pulse height (3,608 pulses). **d,** Cross-correlation between growth rate and *rpoS* activity. **e,** Histogram of growth rate at one frame from all movies for *WT* and *ΔrpoS.* In (**c-e**) the mean ± std dev is plotted with the line and the shaded region, respectively for *WT* (11 technical replicates drawn from 7 biological replicates, 563 mother cells) and *ΔrpoS* (10 tech. reps. drawn from 6 bio. rep., 279 mother cells). **f,** Schematic illustration of mathematical model. Stochastic molecular reactions occur in a growing cell. The reactions are simulated with the Gillespie algorithm, while cell growth happens at deterministic time steps. Growth at each time step is dependent on molecular concentration via Hill functions. The result is a mutual inhibition between growth rate and *rpoS* concentration. **g-j** Analysis from 1,000 simulations run for 500 hours; only the last 250 hours are used to capture steady-state behaviour. **g,** Histograms of simulated *rpoS* concentration with and without feedback of *rpoS* on growth rate (88,865 and 133,126 pulses, respectively). **h,** Cross-correlation between simulated growth rate and *rpoS* concentration. **i,** Histograms of growth rate sampled at 24 hour intervals over all 1,000 simulations to mitigate effects of correlations. **j,** Sample time traces of simulated *rpoS* concentration and elongation rate for four cells. Grey vertical lines indicate cell divisions.

**Figure 3.**
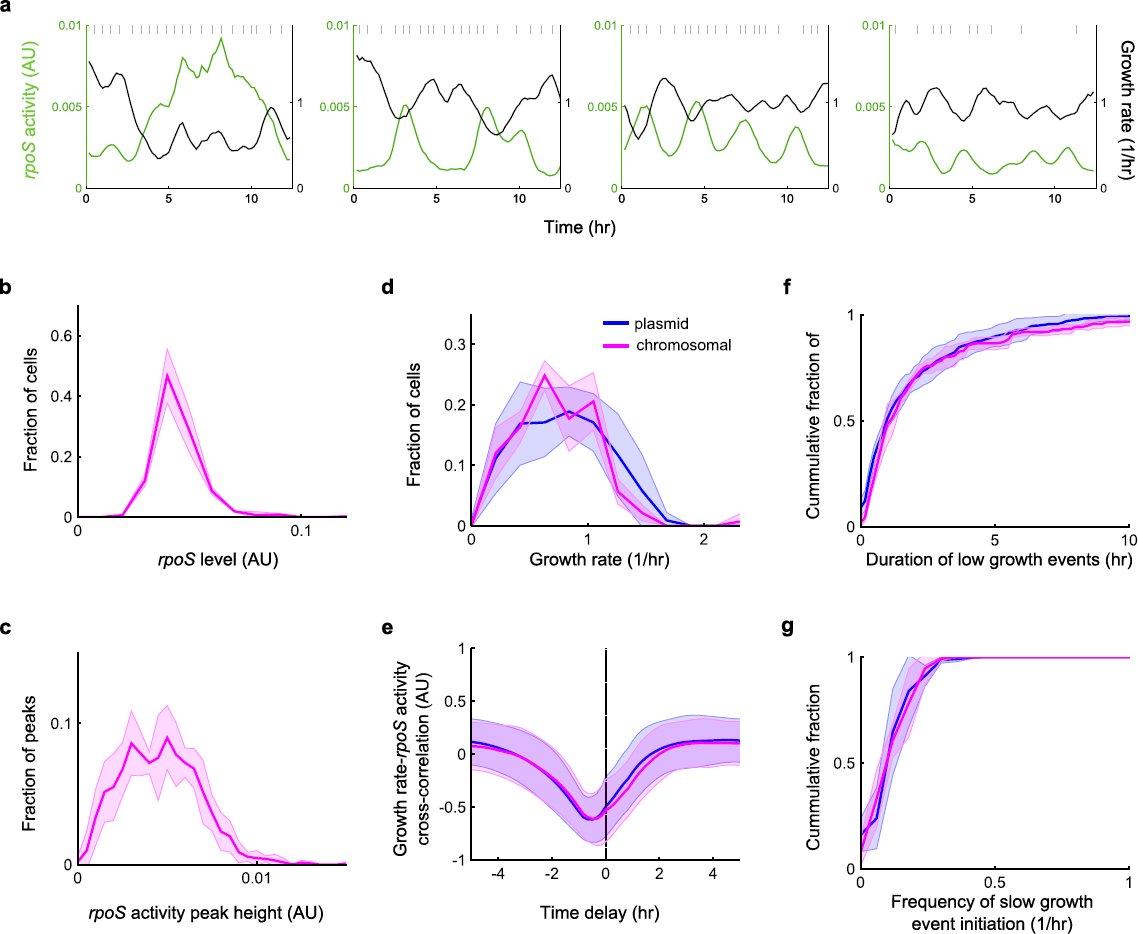
Long-tailed *rpoS* distribution is not due to plasmid segregation effect, nor are the growth effects due to plasmid toxicity. Using chromosomally integrated PboA-GFP in WT: **a,** Sample time traces of *rpoS* activity and growth rate for four mother cells. Grey vertical lines indicate cell divisions. **b,** Distribution of *rpoS* level from bulk liquid culture (3 biological replicates, 465 cells, mean = 0.045 AU, CV = 0.23). **c,** Pulse height distribution in Mother Machine experiments (1,438 peaks). **d,** Growth rate histogram, **e,** Cross-correlation between growth rate and *rpoS* activity. **f,** Duration distribution of low growth events. **g,** Distribution of frequency of entering low growth event. In **c**-**g**, 4 technical replicates drawn from 2 bio. reps., 143 mother cells were used. The plasmid data in **d**-**g** is reproduced from elsewhere in this work for ease of comparison. Lines and shaded region are mean ± std dev, respectively.

We further observed rich dynamics in the growth rate of single cells (Fig. 2b, Methods). The sample lineages illustrate that cell growth slows down when *rpoS* activity is high. This relationship was quantified as a large negative value near zero time-shift in the cross-correlation of growth rate and *rpoS* activity (Fig. 2d, Sup. Fig. 3e, Methods). The strong anti-correlation suggested that growth rate should also be widely distributed, which is what we observed (Fig. 2e, Sup. Fig. 3d, 4b). However, the *ΔrpoS* strain also had a wide growth rate distribution suggesting growth rate is intrinsically heterogeneous^3^ (Fig. 2e). Furthermore, *σ*^70^ activity was positively correlated with growth rate suggesting it is related to this intrinsic variability (Sup. Fig. 4a).

**Figure 4.**
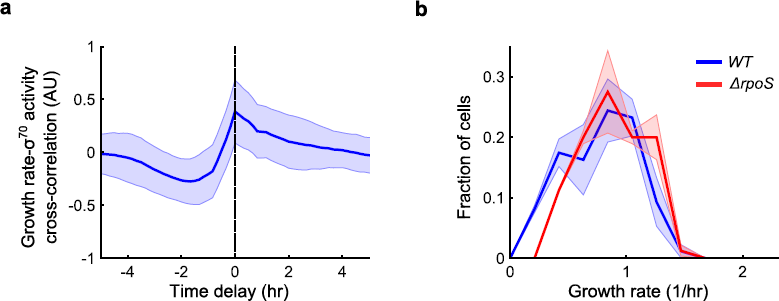
Constitutive, *δ^70^,* reporter is positively correlated with growth and high GFP expression does not affect growth rate distribution. Using *P_rpsL_-GFP* in *WT* and *ΔrpoS.* **a,** Cross-correlation between growth rate and *δ^70^* activity in WT cells. **b,** Growth rate histogram for *WT* and *ΔrpoS. WT:* 2 biological replicates, 86 mother cells; *ΔrpoS:* 2 biological replicates, 81 mother cells. Lines and shaded region are mean ± std dev, respectively.

We propose a coupled molecular and physiological model to explain our observations. First, we propose the intrinsic variability in growth rate arises due to stochastic molecular reactions that promote growth. Second, *rpoS* molecules repress growth and growth dilutes *rpoS.* This results in the anti-correlation between growth rate and *rpoS.*

To test our proposal we constructed a mathematical model. For simplicity, we chose to model two molecular species, growth factor *(γ)* and *rpoS (r).* We used a stochastic Gillespie simulation for the reactions. Both were assumed to be produced by zeroth order reactions and degraded by first order reactions (Fig. 2f, see Methods for details). The reactions occurred in a cell which grew at deterministic time intervals. As the cell volume increased molecule concentration was diluted. The growth rate at each deterministic time step explicitly depended on the most recent *γ* and *rpoS* concentration via the product of Hill functions (Fig. 2f). The Hill function for *γ* rose with concentration while that for *rpoS* decreased. This captured the promoting and repressing effects on growth rate of the two kinds of molecules, respectively.

This coupled molecular and physiological simulation can be summarized as a mutual inhibition feedback between *rpoS* and growth rate^19^ (Fig. 2f). Using a coarse-grained exploration of the parameter space we found parameters for the stochastic simulation and Hill functions which reproduced the *WT* and rpoS-knockout experimental growth distributions (Fig. 2i) as well as the population growth rate. With these parameters set, the model then produced a long-tailed distribution of *rpoS* pulse heights, which decreased in prominence when the negative *rpoS* feedback on growth rate was removed *in silico* (Fig. 2g). The model also captured the rich single-cell *rpoS* and growth dynamics observed (Fig. 2j), as well as the anti-correlation between growth rate and *rpoS* (Fig. 2h).

We tested our understanding of the feedback model by perturbing population growth rate. As population growth rate is reduced, *rpoS* levels should increase due to decreased dilution (Fig. 3a). We reduced population growth rate by reducing culture temperature, using reduced quality media, or combinations of the two (see Sup. Tab. 2) and imaged single cells from bulk cultures (see Methods). Indeed, *rpoS* levels increased with decreasing population growth rate (Fig. 3b).

The ability of *rpoS* to reduce growth rate could decrease with population growth rate due to globally reduced rates of transcription^20,21^. On the other hand, *rpoS* efficacy could remain constant, or even increase, allowing *rpoS* to control a greater portion of transcription. We used the model to distinguish between these possibilities. We modelled a reduction in population growth rate by decreasing *g_max_* (see Methods). The effect of *rpoS* on growth rate could scale with this maximum growth rate, reflecting a constant *rpoS* efficacy, or remain fixed, reflecting an attenuated *rpoS* efficacy. We modelled the former by keeping *ƒ* constant in the *rpoS* Hill function as *g_max_* was varied. The latter was done by keeping the product *fg_max_* constant, thereby flattening the repressive Hill function (Fig. 3c, Methods).

Comparing the theory to experiments, we found *rpoS* efficacy reduced with population growth rate. Using the Mother Machine assay and reduced culture temperatures we experimentally observed a convergence of the growth rate distributions of *WJ* and *ArpoS* populations (Fig. 3e, Sup. Fig. 5b). We found that the constant efficacy model overestimated the effect of *rpoS* on single-cell growth rate as population growth rate was reduced (Fig. 3d, e Sup. Fig. 5a, b), whereas the reduced *rpoS* efficacy model faithfully represented reality (Fig. 3e, f, Sup. Fig. 5b, c). Additionally, the reduced efficacy model captured the increasing levels of *rpoS* at reduced population growth rates (Fig. 3b).

**Figure 3.**
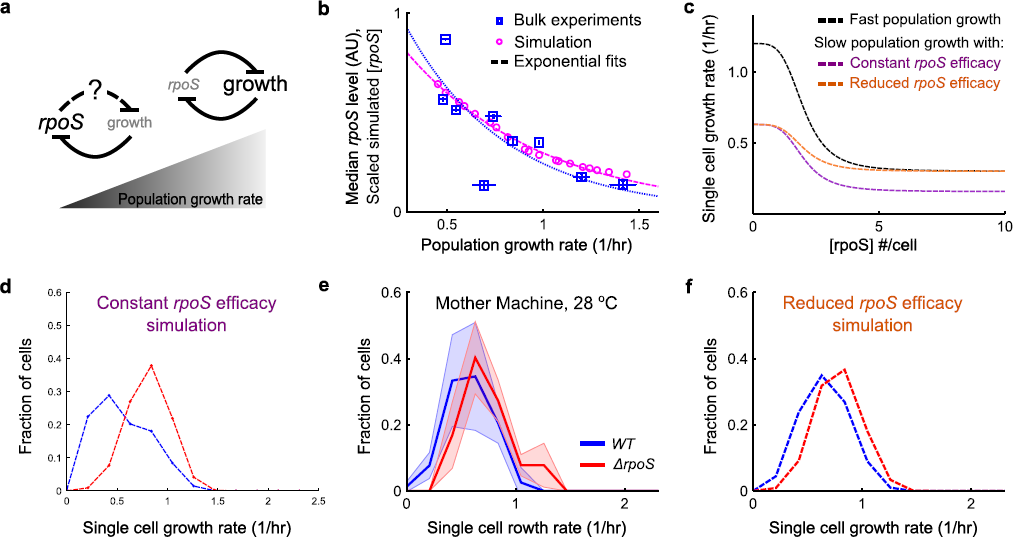
*rpoS* levels increase, but are less potent, at reduced population growth rate. **a,** Schematic illustrating effect of reduced population growth rate. *rpoS* is concentrated due to lower dilution by growth rate. However, its effect on growth rate could diminish at low population growth rate. **b,** Median *rpoS* levels in liquid culture (± std dev, mean growth rate ± std dev, at least two biological replicates, see Sup. Tab. 2 for details) and scaled *rpoS* concentration from simulations as functions of population growth rate. Dashed lines are exponential fits. Scaling factor (0.29) was found by minimizing root-mean-square error between the fits over the range of observed growth rates ± 20% (0.29 to 1.6/hr). **c,** Hill functions of growth rate as functions of *rpoS* concentration used in simulations. Fast population growth corresponds to simulation matching experimentally observed growth rate at 37°C (Fig. 2e, i). The constant and reduced efficacy models behave differently in the large *rpoS* concentration limit as population growth rate is reduced. **d-f,** Growth rate histograms for *WT* and *ΔrpoS.* **e**, Cells grown at reduced temperature, 28°C, in the Mother Machine (mean ± std dev, WT, 4 technical replicates drawn from 3 biological replicates, 84 mother cells; *ΔrpoS,* 4 tech. reps. drawn from 2 bio. rep., 85 mother cells). Simulation results at corresponding population growth rate with constant *rpoS* efficacy (**d**) and reduced *rpoS* efficacy (**f**) (100 simulations for 19 values of *g_max_,* sampled every 24 hours, in the final 250 hours of 500 hour simulations).

The *rpoS* regulon allows cells to survive a variety of environmental stresses, for instance oxidative stress^8,16,22^. To test the function of heterogeneous *rpoS* expression, we assayed the survival of exponential phase cells against hydrogen peroxide (H_2_O_2_). We used a short, intense pulse of stress to study the effect of *rpoS* already present in the bacteria, as opposed to the well-studied stress-induced *rpoS* response^16^. Using the Mother Machine we allowed cells to grow in fresh media, briefly switched to media containing H_2_O_2_, and then back to fresh media (Fig. 4a, see Methods for details). The population of cells that survived the stress upregulated *rpoS* approximately three hours prior to the stress (Fig. 4b). Consistent with literature^16^, *rpoS* knockout populations had a reduced survival fraction compared to *WT* (Fig. 4f).

**Figure 4.**
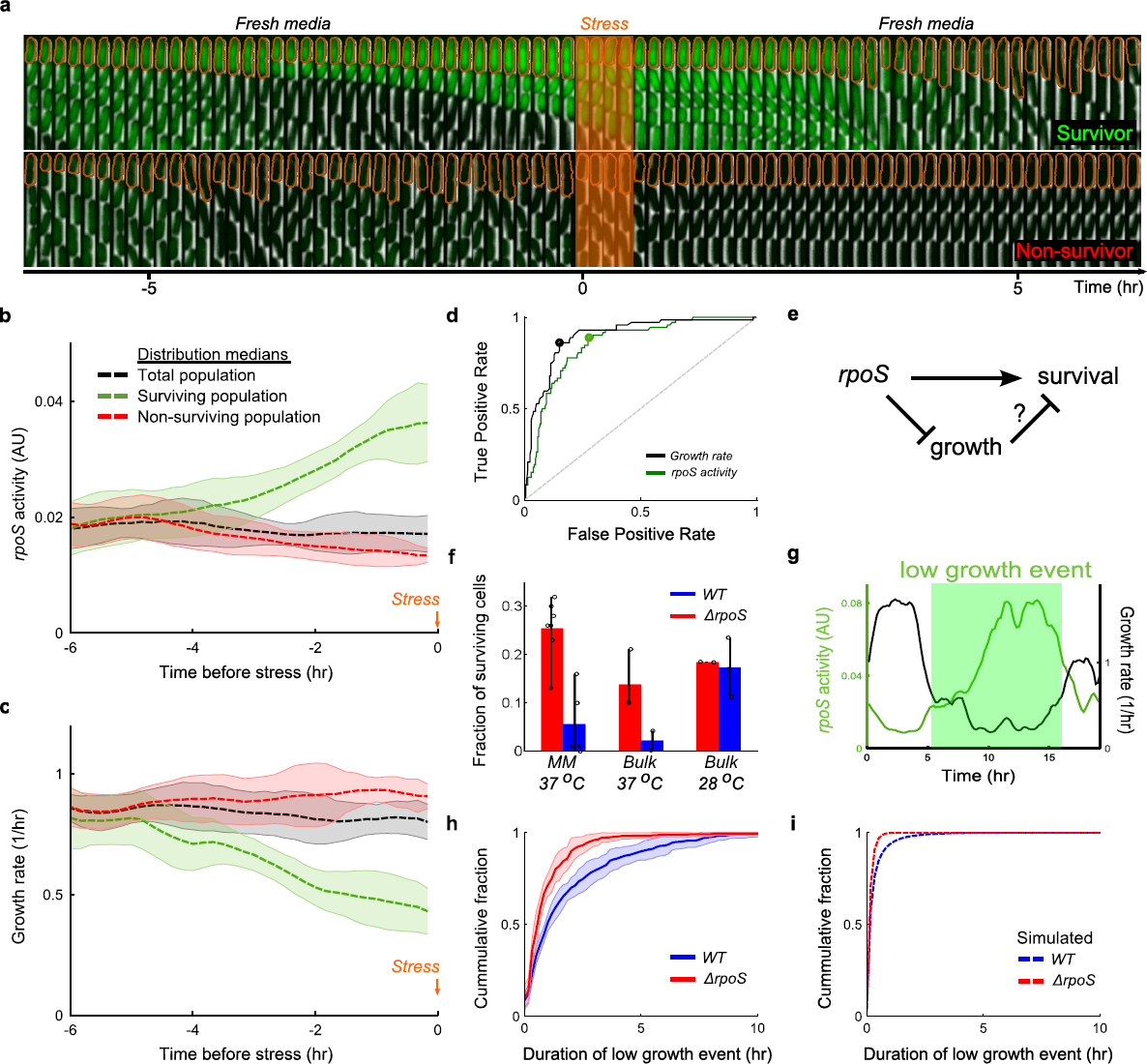
*rpoS* enables survival of stress by prolonging duration of slow growing state. **a,** Schematic of the stress assay and sample montages of surviving (top) and non-surviving (bottom) mother cell. Mother cell outlined in orange; phase contrast and fluorescence channel ranges identical for both montages and chosen for display. Cells were grown for 10 hours in fresh media, followed by a 35 minute application of H_2_O_2_ stress, and fresh media once again. **b**, Median value of *rpoS* activity distributions for time points prior to stress application (t = 0), sorted according to survival (line and shaded area are mean ± std dev, 7 technical replicates drawn from 4 biological replicates; 72 surviving cells, 212 non-surviving, 284 total mother cells). **c**, Same as (**b**) for growth rate. **d,** Receiver Operating Characteristic curve for growth rate (black) and *rpoS* activity (green) from time point preceding stress application. Grey dashed line is True Positive Rate = False Positive Rate. Circles represent locations of optimal thresholds (0.65/hr for growth rate, 0.020 AU for *rpoS* activity). Area Under the Curve (AUC) is 0.90 for growth rate and 0.86 for *rpoS* activity. **e,** Schematic illustrating alternative mechanisms of stress survival. High *rpoS* activity could directly allow cells to survive or it might first reduce growth rate, which in turn allows survival. **f,** Fraction of cells surviving stress in the Mother Machine assay (mean ± min/max, WT: 7 tech. reps., represented as circles, drawn from 4 bio. reps., 1,087 cells, *ΔrpoS:* 5 tech. reps. drawn from 3 bio. reps., 996 cells) and bulk Colony Forming Units assay at two temperatures (mean ± min/max; at least two biological replicates for bulk culture assays, represented as circles). **g,** Illustration of a low growth event based on the ROC curve optimal threshold (0.65/hr) (**d**). **h,** Cumulative distribution of duration of low growth events in *WT* and *ΔrpoS* populations (line and shaded area are mean ± std dev, WT, 11 tech. reps. drawn from 7 bio. reps., 563 mother cells, 941 events; *ΔrpoS,* 10 tech. reps. drawn from 6 bio. rep., 279 mother cells, 391 events). **i,** Same as (**h**) from simulations (1,000 simulations run for 500 hours, only the final 250 hours were used; *WT,* 75,787 events and *ΔrpoS,* 49,114 events).

Intriguingly, the surviving population also had reduced growth rate prior to the stress (Fig. 4c). Using the Receiver Operating Characteristic (ROC) curve we found that both *rpoS* activity and growth rate immediately preceding stress application are strong predictors of survival (Fig. 4d and Methods). This suggested two alternative hypotheses; either *rpoS* directly causes the survival phenotype, or it acts by first reducing growth rate, which in turn allows cells to survive the stress (Fig. 4e).

To distinguish between the two hypotheses, we noted the fraction of cells growing slower than the optimal threshold for survival increased for both *WT* and *ΔrpoS* populations as population growth rate decreased (Fig. 4d, Methods, Sup. Fig. 5b). If *rpoS* directly caused survival, then the difference in survival fraction between *WT* and *#x394;rpoS* populations should increase at reduced temperature due to the increased *rpoS* present in *WT* cells (Fig. 3b). On the other hand, if growth rate was causing survival, the difference should decrease. We tested this experimentally by a bulk culture Colony Forming Units (CFU) stress assay (see Methods) and found the latter (Fig. 4f). Furthermore, we observed rpoS-knockout cells that survived in the Mother Machine assay at 37°C also down-regulated growth prior to stress (Sup. Fig. 6). This prompted the question: What is the role of *rpoS* at fast population growth rates?

**Figure 5.**
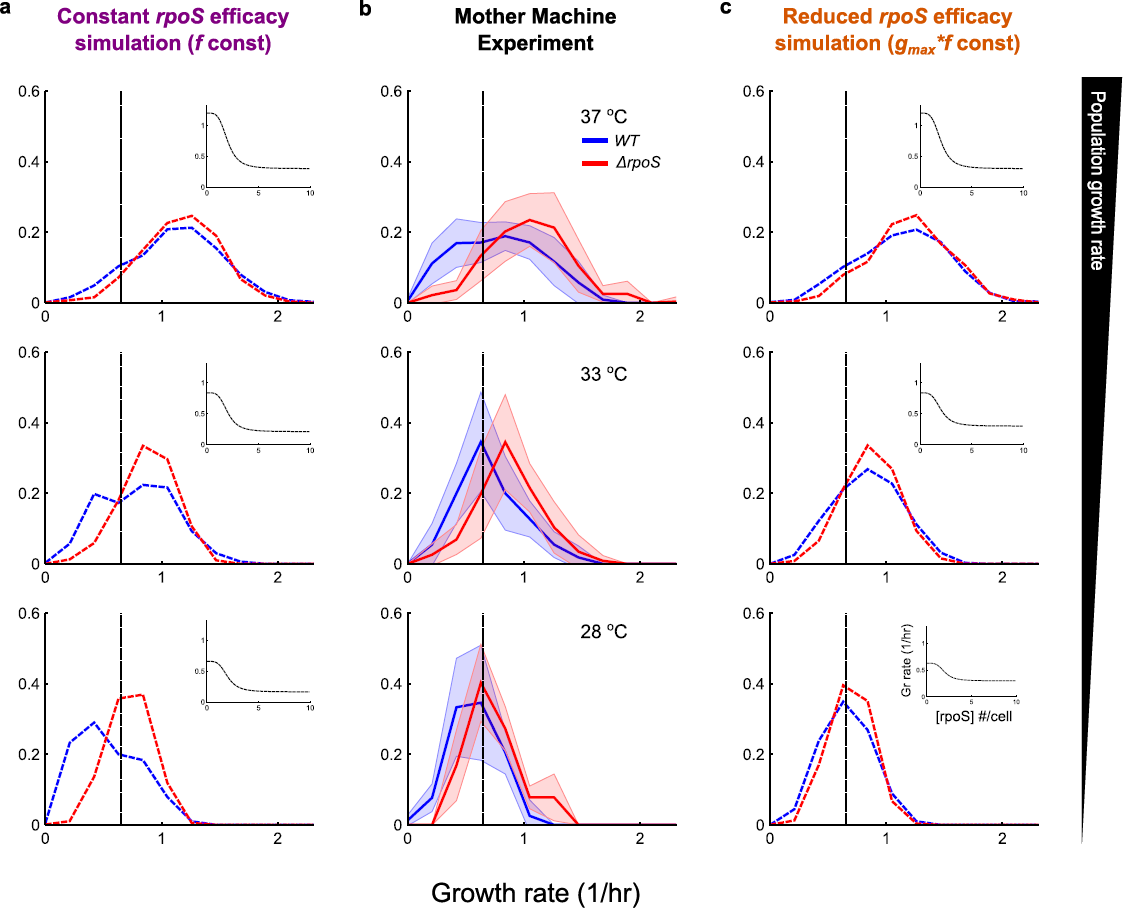
The influence of *rpoS* on growth is attenuated as population growth rate decreases. **a,** Growth rate histograms for simulated *WT* and *ΔrpoS* at three population growth rates achieved by keeping *ƒ* constant as *g_max_* was reduced. Dashed black lines correspond to optimal survival threshold of 0.65/hr (Fig. 4d). Insets: Hill functions of growth rate vs *rpoS* concentration. **b,** Experimental growth rate histograms for *WT* and *ΔrpoS* grown at three temperatures (mean ± std dev, 28°C and 37°C reproduced from main text; 33°C WT, 5 technical replicates drawn from 3 biological replicates, 72 mother cells; *ΔrpoS,* 6 tech. reps. drawn from 3 bio. rep., 137 mother cells). **c**, Growth rate histograms for simulated *WT* and *ΔrpoS* with *F∗g_max_* constant as *g_max_* was reduced. Insets: same as (**a**). *g_max_* values for the simulations were chosen such that population growth rates matched the experimentally observed population growth rates, see Methods for details. For (**a**) and (**c**) 100 simulations were used for each condition, sampled every 24 hours, in the final 250 hours of 500 hour simulations.

**Figure 6.**
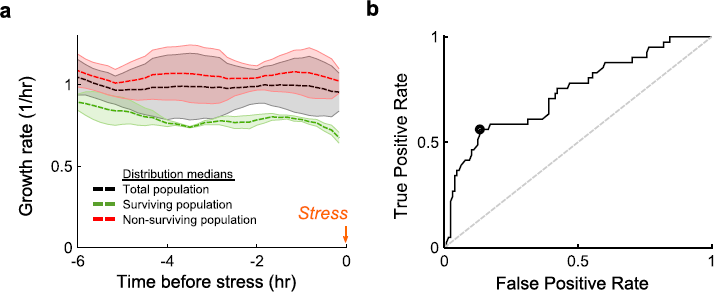
Slow growing *ΔrpoS* cells survive oxidative stress. Cells were treated as in Fig. 3. **a,** Median value of growth rate distributions for time points prior to stress application (t = 0), sorted according to survival (mean ± std dev, 5 technical replicates drawn from 3 biological replicates, 41 surviving cells, 128 non-surviving cells, 169 total mother cells). **b**, Receiver Operating Characteristic curve for growth rate (optimal threshold is 0.70/hr, Area Under Curve is 0.74).

To answer this question we analysed periods when cells were growing slower than the optimal threshold for survival (Fig. 4g). The role of *rpoS* is to prolong the duration of these slow growth events. We observed this as a higher frequency of long duration slow growth events in *WT* compared to *rΔpoS* (Fig. 4h, Sup. Fig. 3f). The frequency with which cells attempt to grow slowly for any duration is similar for *WT* and *ΔrpoS* populations (Sup. Fig. 7a, 3g). The rpoS-growth feedback model captures this dynamic *rpoS* phenotype (Fig. 4i, Sup. Fig. 7b).

**Figure 7.**
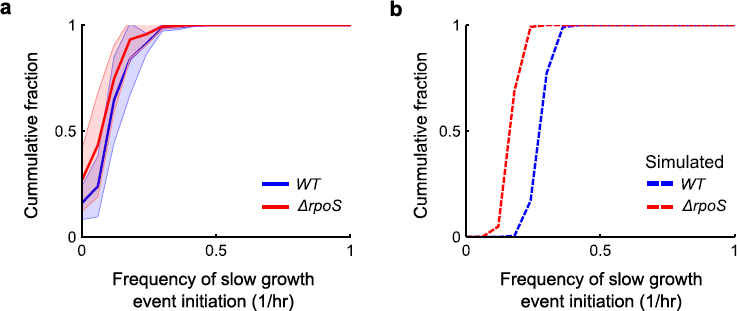
Frequency of slow growth initiation similar between WT and rΔpoS. **a,** Experimental distributions of frequency of entering low growth event for *WT* and *ΔrpoS* (mean ± std dev; WT, 11 technical replicates drawn from 7 biological replicates, 563 mother cells, 821 events; *ΔrpoS,* 10 tech. reps. drawn from 6 bio. rep., 279 mother cells, 342 events). **b,** Same as (**a**) for simulations (1,000 simulations run for 500 hours, only the final 250 hours were used; *WT,* 75,628 events and *ΔrpoS*, 49,041 events).

Slow growth mediated by *rpoS* has been implicated in the closely related persistence phenomenon^23^. Persister cells are slow growing cells in a clonal population of otherwise fast growing cells that can survive transient antibiotic treatment^5,6^. However, sudden downshifts in nutrient quality can generate nearly homogenous persister populations that are characterised by upregulation of the *rpoS* regulon^23^. We wondered if the heterogeneously generated surviving cells we observed were in fact persisters. There are several key differences. First, cells surviving oxidative stress were able to grow ~30x faster than persisters^6,23^. These survivors also occurred several orders of magnitude more frequently^6^. Finally, the small molecule ppGpp has been implicated in the production of heterogeneous persister cells^24^. We found that cells devoid of the ppGpp synthase, *relÄ^25^,* exhibited similar long-tailed *rpoS* distributions as wild type cells (Supp. Fig. 8), further distinguishing *rpoS* survivors from persisters.

**Figure 8.**
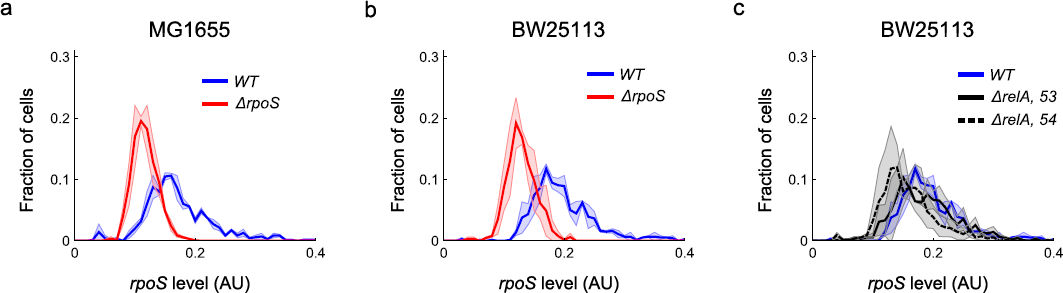
ppGpp does not affect long-tailed *rpoS* expression. **a,** *WT* strain used in this work, MG1655, and *ΔrpoS::kan* harbouring reporter with kanamycin resistance replaced with spectinomycin resistance *(P_bolA_-GFP::spec). WT* (2 biological replicates, 696 cells, mean = 0.18 AU, CV = 0.38) and *ΔrpoS* (2 bio. reps., 1,244 cells, mean = 0.12 AU, CV = 0.18). **b,** The same in the *WT* strain of the Keio collection^28^, BW25113. *WT* (2 bio. reps., 739 cells, mean = 0.20 AU, CV = 0.27) and *ΔrpoS* (2 bio. reps., 651 cells, mean = 0.13 AU, CV = 0.19). **c,** Two isolates from the Keio collection of *ΔrelA,* from plates 53 and 54 with *WT* from (**b**): *ΔrelA, 53* (2 bio. reps., 898 cells, mean = 0.19 AU, CV = 0.29) and *ΔrelÄ, 54* (2 bio. reps., 543 cells, mean = 0.16 AU, CV = 0.29). Lines and shaded region are mean ± std dev, respectively.

Despite these differences, the two phenomena may be connected. Exposure to antibiotics can enhance subsequent survival against acid stress, a response mediated by *rpoS^26^.* Perhaps *E. coli* experienced antibiotic stress simultaneously with harsh environments in its evolutionary history. Cells able to coordinate persistence with the *rpoS* survival strategy revealed here would out-compete uncoordinated cells.

We combined theory and experiments to reveal how mutual inhibition of *rpoS* and growth can generate a rich, dynamic phenotype. Our coupled stochastic molecular and cell growth model provides a platform to explore more detailed mechanistic models. We have also demonstrated how the predictions of such a theory can be fruitfully compared to quantitative single-cell data. The active degradation of *rpoS* by proteases^8^ and the promotion of anti-δ^70^, *rsd,* by *rpoS^27^* are two mechanisms that could provide greater agreement between theory and experiments. We therefore anticipate more functional phenotypic variability will be revealed by this approach.

## Methods

### Strains and growth conditions

#### Media

M9 (1xM9 Salts, 2mM MgSO_4_, 0.1 mM CaCl_2_; 5xM9 Salts 34g/L Na_2_HPO_4_, 15g/L KH_2_PO_4_, 2.5 g/L NaCl, 5 g/L NH_4_Cl) supplemented with 0.2% Casamino acids and 0.4% glucose as carbon source. Media for Mother Machine experiments was also supplemented with 0.2 mg/mL Bovine Serum Albumin (BSA). For growth rate perturbation experiments glucose was replaced with 0.4% mannose and Casamino acids with 1 mM thiamine (see Sup. Tab. 2 for further details).

**Table S.2.** Growth conditions and population growth rates

| Growth condition* | WT |  | *ΔrpoS* |  |
| --- | --- | --- | --- | --- |
|  | Biological replicates; number of cells | Growth rate (1/hr), mean ± std dev | Biological replicates; number of cells | Growth rate (1/hr), mean ± std dev |
| 0.4% glucose, 0.2% casamino acids (37°C) | 4; 711 | 1.42 ± 0.07 | 3; 427 | 1.42 ± 0.08 |
| 0.4% glucose, 0.2% casamino acids (33°C) | 2; 547 | 0.98 ± 0 | 2; 510 | 1.02 ± 0 |
| 0.4% glucose, 0.2% casamino acids (28°C) | 2; 747 | 0.55 ± 0.02 | 2; 601 | 0.59 ± 0.01 |
| 0.4% mannose, 0.2% casamino acids (37°C) | 3; 720 | 1.20 ± 0.04 | 2; 453 | 1.23 ± 0 |
| 0.4% mannose, 0.2% casamino acids (33°C) | 2; 346 | 0.84 ± 0 | 2; 511 | 0.85 ± 0.02 |
| 0.4% mannose, 0.2% casamino acids (28°C) | 2; 604 | 0.48 ± 0.02 | 2; 595 | 0.51 ± 0.01 |
| 0.4% glucose, 1 mM thiamine (37°C) | 2; 896 | 0.74 ± 0.04 | 2; 536 | 0.74 ± 0.02 |
| 0.4% mannose, 1 mM thiamine (37°C) | 3; 2,719 | 0.49 ± 0.02 | 3; 2,298 | 0.52 ± 0.03 |

\*M9 supplemented with the following and grown at (temperature).

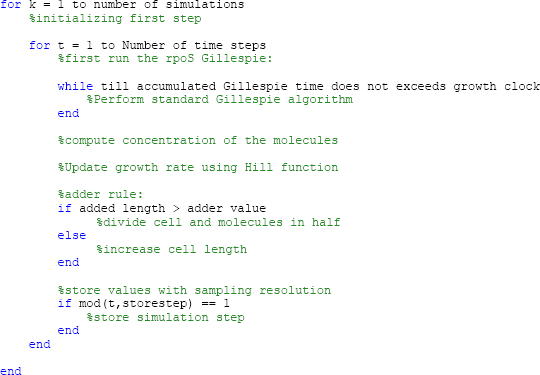
*Pseudo code*

*Reporter plasmid*

Reporter plasmids were sourced from the Alon library^18^ using standard procedures and Qiagen Miniprep kits. Strains were transformed with the appropriate reporter plasmids by using a variant of the Top10 Chemical Competence protocol (OpenWetWare) followed by standard transformation by heat shock. Either an overnight culture or cells taken directly from glycerol stocks were grown up to exponential phase in LB. The cells were washed and concentrated in pre-chilled CCMB80 buffer 2-3 times (CCMB80: 10mM KOAc, 80 mM CaCl_2_-2H_2_0, 20 mM MnCl_2_-4H_2_O, 10 mM MgCl_2_-6H_2_O, 10% glycerol, adjusted to pH 6.4 with HCl). Next the plasmid was added to the cells and the mixture incubated on ice for 20-30 minutes. After a 1 minute 42°C heat shock, cells were allowed to recover in 1 mL LB at 37°C for 1 hour before plating on LB agar plates with 25ug/mL Kanamycin selection overnight. See Sup. Tab. 1 for list of strains.

**Table S.1.** Strain list

|  | Strain name | Genotype | Construction procedure | Source |
| --- | --- | --- | --- | --- |
| 1a | MG1655 seq (CGSC#6300) | K12 with *F-λ-rph-1* | WT strain of reporter library<sup>19</sup> | Yale CGSC |
| 1b | MG1655 (CGSC#7740) | K12 with *F-λ-rph-1* | WT strain of reporter library<sup>19</sup> | Yale CGSC |
| 2 | 63Dr | Same as 1a with *ΔrpoS::Kan<sup>r</sup>* | Used a PCR product from Keio collection *ΔrpoS* strain<sup>29</sup> | This work |
| 3a | 63DrF- | Same as 2, markerless | FLPe recombinase | This work |
| 3b | MGDrF- | Same as 1b with *ΔrpoS* | Used a PCR product from Keio collection *ΔrpoS* strain<sup>29</sup> and FLPe recombinase | This work |
| 4 | MGChrRep | Same as MG1655 with chromosomally integrated reporter::*kan* | Used Red/ET system and PCR product amplified from reporter plasmid<sup>19</sup> | This work |
| 4 | BW25113 | Same as MG1655 with *rrnB3 ΔlacZ4787 hsdR514 Δ(araBAD)567 Δ(rhaBAD)568* and *Δcrl ΔvalX mhpC365991* | WT strain of Keio collection<sup>29</sup> | Yale CGSC |
| 5 | KDr | Same as BW25113 with *ΔrpoS::Kanr* | - | From Keio collection<sup>29</sup> |
| 6a | DrelA53 | Same as BW25113 with *ΔrelA::Kanr* | - | From Keio collection<sup>29</sup> (plate 53) |
| 6b | DrelA54 | Same as BW25113 with *ΔrelA::Kanr* | - | From Keio collection<sup>29</sup> (plate 54) |

#### Knockout construction

Knockouts strains were sourced from the Keio collection^28^. The knockout site with Kan^r^ was amplified by PCR and used to perform knockouts in the MG1655 *E. coli* strain. Knockouts were carried out by the commercial Red/ET Recombination system (Gene Bridges, Germany) following the recommended protocol. However, instead of electroporation for transforming with the Red/ET recombination plasmid and FLPe flipase plasmid we used chemical transformation. The transformation was as above except the recovery was carried out at 30°C and 1,000 rpm in a benchtop shaker and plates incubated at 30°C as the plasmid replication ceases at 37°C. Knockouts were verified by colony PCR and sequencing.

#### Chromosomal integration of reporter

Knockins were performed as above for knockouts with the Red/ET recombination system (Gene Bridges). The integrated DNA was amplified off the reporter plasmid. The reporter plasmids were sequenced and used as references for the integration.

#### Mother Machine microfluidic device

The Mother Machine microfluidics device has been described previously^10^. Briefly, it consists of a feed trench (~50 µm x 100 µm x 30 mm) with many channels (~1.4µm x 1.4 µm x 25µ m) attached perpendicular to the trench. These channels hold the cells and media is supplied to the cells via the trench. We used an epoxy master mould to fabricate our devices, which was a kind gift of Suckjoon Jun. The devices were fabricated by casting Sylgard 184 polydimethylsiloxane (PDMS) (Dow Corning, USA) with a ratio of 10:1 base to curing agent onto the master mould and cured overnight at 65°C. The chips were then cut out and plasma bonded (Femto Plasma System, Diener, Germany) to a glass bottom dish (HBSt-5040, Wilco Wells, Netherlands). To strengthen the bonding the chips were incubated for approximately ten min at 65°C. The chips were passivated with 20 mg/ml Bovine Serum Albumin (BSA) for approximately one hour at 37°C prior to cell loading.

### Data acquisition

#### Bulk culture snaps

We used the imaging protocol described previously^9^ with minor modifications. Cells were grown from glycerol stocks in M9 at 37°C to late exponential phase and then diluted back into M9 to an OD of 0.01. After regrowing for approximately 2 hours 20 minutes, up to early exponential phase (OD~0.15), 0.3 µL of the cell culture was spotted onto pads of 1.5% low-melting agarose in Phosphate-Buffered Saline (PBS). Cells were imaged expediently, typically within ~20 minutes of leaving the incubator.

#### Population growth rate perturbation

Cells were grown from glycerol stocks using the modified media and temperature into exponential phase. Optical density measurements were taken after cells were diluted and grown up to exponential phase for imaging.

#### Mother Machine movies

Cells were grown from glycerol stocks as above. They were concentrated by centrifugation (4,000 rpm for 10 min) and injected into the Mother Machine devices. A second centrifugation step for 5 min at 4,000 rpm using a spin coater (Polos Spin 150i, SPS, Netherlands) forced cells into the channels. Cells were allowed to settle in the device while being supplied with fresh media for ~2 hours prior to beginning acquisition. Media was supplied at a flowrate of 1 ml/h by either a Fluigent pressure pump (MFCS-EZ, Fluigent, France) with an M-Flow sensor (Fluigent, France) or a syringe pump (Fusion 100, Chemyx, USA).

#### Microscopy

We used a widefield microscope with epifluorescence and phase contrast imaging modes (Nikon Ti-eclipse, Nikon, UK) equipped with the Nikon Perfect Focus (PFS) Unit. Illumination for the epifluorescence was provided by a white light LED source (SOLA SE Light Engine or Spectra X Light Engine, Lumencor, USA), transmitted by a liquid light guide (Lumencor, USA), through a fluorescence filter cube (49002-ET-EGFP, excitation: ET470/40x, dichroic: T495LP, emitter: ET525/50m, Chroma, USA), and a CFIPlan Apochromat 100x oil immersion objective (NA 1.45, Nikon). Phase contrast illumination was provided by a 100 W lamp via a condenser unit (Nikon). Images were acquired on a CoolSNAP HQ^2^ camera (Photometrics, USA). The sample was held in motorized stages (Nikon). The sample was incubated along with much of the microscope body using a temperature controlled, heated chamber (Solent Scientific, UK). The microscope was controlled with MetaMorph software (version 7.8.10.0, Molecular Devices, USA). Fluorescent beads (TetraSpeck microspheres, 0.5 µm, Molecular Probes, USA) were imaged as a calibration standard.

### Quantifying gene expression and growth rate

#### Bulk culture single-cell gene expression

A custom MATLAB (Mathworks, USA) script based on the published Schnitzcells software was used for image analysis^9^. The microscope was calibrated for each experiment with fluorescent beads to mitigate the effect of non-uniform sample illumination and daily variations in the apparatus. Cells were taken from a field of view computed from the beads to be within 80% of maximum intensity. Cells were segmented in the phase contrast channel. The mean fluorescence was then the corresponding pixels in the GFP channel normalized to cell area. A threshold was applied to exclude debris and substrate autofluorescence was subtracted from the mean cell fluorescence. Finally, the cell fluorescence was normalized by the fluorescence of the top 2% of fluorescent beads.

#### Growth perturbation experiments

Gene expression was computed as above. Growth rate was calculated by fitting an exponential curve to the OD measurements.

#### Mother Machine movies

Cell segmentation was done on the phase contrast channel. The mother cell - the cell that remained at the end of growth channels farthest from the feed trench - was isolated and tracked. The image analysis was robust most of the time, but failed intermittently. Thus, every frame used in subsequent analysis was manually checked, and corrected if necessary. Growth was assumed to be exponential for each cell^29^, *i.e. dl/dt = g*∗*l*, where *l* is cell length and *g* the growth rate. We thus computed growth rate as the difference in cell length between consecutive frames normalized by the first length. We note that the Mother Machine technique overrepresents slow growing cells compared to bulk culture since the slow growing cells do not have to compete with fast cells in the Mother Machine. The population growth rate of mother cells was computed as *g_pop_* = *ln(2)/t_D_* where *t_D_* was found by numerically solving:

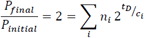

Where *P_x_* are number of cells, *n_i_* are the fraction of cells growing with cell cycle time *c.*

#### Promoter (rpoS) Activity

Gene expression level was calculated as above. Calibration to beads was done using only the top 2% normalization - no cells were excluded due to position in the field of view. Promoter activity (A) was computed as the component of the time-derivative of the expression corrected for by growth rate and bleaching^1^:

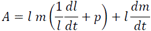

Where *I* is cell length as above, *m* is mean fluorescence, and *p* is an adjustable parameter accounting for photobleaching of GFP. We set *p* = 0.1.

#### Cross-correlation

The normalized cross-correlation between growth rate and promoter activity was computed as follows:

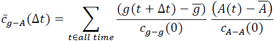

Where *g* is growth rate, *A* is promoter activity, Δt, is the time difference between the two signals, overbars indicate averages over time, and *c* is the auto-correlation:

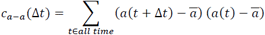

Where a is either promoter activity or growth rate.

### Survival assay

#### Mother Machine assay

Cells were loaded into the Mother Machine as above. Cells were allowed to grow in fresh media for 10 hours, then exposed to 35 mM H_2_O_2_ for 35 minutes and then supplied with fresh media again for at least 12 hours. The media was switched with a Fluigent 2-switch or M-switch (Fluigent, France). Two 35 minute pulses of 3 to 12 mM propidium iodide were supplied with the second round of fresh media and the cells were imaged in the RFP channel to observe DNA chelation of dead cells. This approach was not robust for identifying survivors and dead cells. Thus the movies for each mother cell were manually curated to determine survival using solely the phase contrast channel. If the cell began growing post-H_2_O_2_ treatment and before the movie ended, it was counted as a survivor. Ambiguous cases were excluded from the tally *(WT,* 14% of cells excluded, *ΔrpoS,* 5%), however including these cells in the survival fraction calculation did not change the results.

#### Receiver Operating Characteristic (ROC) curve

A ROC curve measures how well a binary classifier performs as the threshold of the classifier is varied. We used growth rate and *rpoS* activity to classify the survival of cells in the Mother Machine survival assay. The True Positive Rate (TPR) as a function of the threshold was computed as:

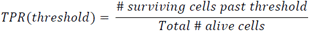

Similarly, the False Positive Rate (FPR) was computed as:

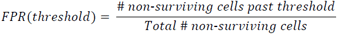

When growth rate was used as the classifier, cells passed the threshold if their growth rate was below the tested value; while for *rpoS* activity if it was above. The TPR was plotted against the FPR to generate the ROC curve. The optimal threshold was computed by finding the threshold that resulted in the maximum difference between the TPR and FPR. The Area Under the Curve (AUC), computed by numerical integration of the ROC curve, is a measure of the quality of the classifier. A perfect classifier has AUC = 1, while one that is no better than random guessing has AUC = 0.5.

#### Bulk culture Colony Forming Units (CRÛ) assay

Cells were grown into exponential phase from glycerol stocks at either 37°C or 28°C and diluted into 10 mL fresh media. They were grown into exponential phase again and aliquoted into 2 mL cultures. These aliquots were exposed to either water or 26 mM H_2_O_2_ and incubated for a further 20 minutes. Cultures were then serially diluted in M9 and plated on LB agar plates. The colonies on the plates were counted after an overnight incubation at 37°C to determine the Colony Forming Units (CFU). Survival fraction was computed as cells/mL from the stress condition divided by the cells/mL from the water condition. Averages were taken over all plates that were in the dynamic range of the assay (30 to 300 colonies per plate).

### Stochastic simulation coupled to single cell growth model

We modelled a single cell growing as a function of molecular reactions occurring inside it. A single lineage was followed, *i.e.* only one daughter cell was followed at each cell division. To model growth, we assumed rodshaped cells with fixed radius and modelled growing cells by the changing length at a fixed, deterministic time interval, *Δt*:

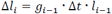

where *g_i_* and *l_i_* are the growth rate and cell length at the *i^th^* time point, respectively. Cell division was assumed to follow the adder rule^29^:

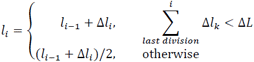

where *ΔL* is a fixed length the cell must add before it can divide. The numbers of molecules in the cell were determined by a standard Gillespie stochastic simulation algorithm^11^ that ran between the deterministic steps of the growth model. Two molecular species *rpoS, r,* and growth factor, *γ,* were modelled. They were generated with zeroth order constitutive production and first order degradation reactions:

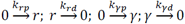

where *k_xp_* are the production propensities and *k_xd_* are the degradation propensities for species ***x***. The reaction propensities in the Gillespie algorithm do not change with cell volume since the reactions are zeroth and first order^12^. At division the number of molecules were simply divided in half and rounded to the closest integer lower than the quotient:

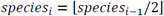

The concentration of the molecular species was the number of species divided by cell length (volume):

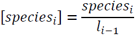

Growth rate was a function of the concentration of the two molecular species generated most recently by the Gillespie algorithm:

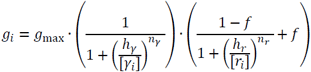

where *g_max_* is the maximum growth rate; *f* represents the lowest growth rate can be reduced to in the limit of infinite *rpoS* concentration; *h_γ_* and *h_γ_* are the values of growth factor and *rpoS* leading to half-maximal growth, respectively; and *n_γ_ n_r_* are the Hill coefficients. Growth factor was considered a downstream target of *δ^70^* so *n_γ_* was positive, while *n_r_* was chosen to be negative to capture the repressive effect of *rpoS* on growth. Growth perturbation simulations were implemented by varying *g_mcm_* all other parameters were kept constant. However, in the reduced *rpoS* efficacy model the parameter *f* was increased to keep the product *fg_max_* constant. See Sup. Table 3 for parameter values used and Sup. Mat. for the pseudo code of the algorithm.

**Table S.3.** Model Parameters

| Parameter | Value in model | Value in physical units | Description |
| --- | --- | --- | --- |
| *Gillespie* |  |  |  |
| $k_{\gamma p}$ | 2.2 | $13 \text{ hr}^{-1}$ | $\gamma$ zeroth order production rate constant |
| $k_{\gamma d}$ | 0.2 | $1.2 \text{ hr}^{-1} \gamma^{-1}$ | $\gamma$ first order degradation rate constant |
| $k_{rp}$ | 0.3 | $1.8 \text{ hr}^{-1}$ | *rpoS* zeroth order production rate constant |
| $k_{rd}$ | 0.01 | $0.06 \text{ hr}^{-1} r^{-1}$ | *rpoS* first order degradation rate constant |
| $\gamma_{init}$ | 1 | 1 molecule | Initial value of $\gamma$ |
| $r_{init}$ | 1 | 1 molecule | Initial value of *rpoS* |
| *Growth* |  |  |  |
| $\Delta L$ | 1 | 2 $\mu\text{m}$ | Length cell must grow before dividing |
| $l_i$ | 1 | 2 $\mu\text{m}$ | Initial cell length |
| *Coupling growth and Gillespie models* |  |  |  |
| $g_{max}$ | 1.2 | $7.2 \text{ hr}^{-1}$ (0.7 $\text{hr}^{-1}$) | Maximum (average) growth rate achievable by cell |
| $h_{\gamma}$ | 17 | 17 molecules/cell | Half-maximum value for γ-growth Hill function |
| $n_{\gamma}$ | 1 | - | Hill coefficient for γ-growth Hill function |
| $h_r$ | 2 | 2 molecules/cell | Half-maximum value for *rpoS*-growth Hill function |
| $n_r$ | -4 | - | Hill coefficient for *rpoS*-growth Hill function |
| $f$ | 0.25 | - | Minimum value *rpoS* can reduce growth rate by |
| *Technical parameters* |  |  |  |
|  | 100 or 1,000 | - | Number of simulations |
|  | 3000 | 500 hrs | Number of time steps |
|  | 0.005 | 3 s | Simulation time resolution |
| | $1/0.005 = 200$ | 10 min | Sampling resolution |

### Code availability

Code used for simulations and for analysis of data reported in this study is available upon request from the corresponding author.

### Data availability

Data that support the findings reported in this study are available upon request from the corresponding author.

## Acknowledgements

We thank Casandra Villava for assistance with cloning; JT Saul and Suckjoon Jun for the kind gift of a Mother Machine epoxy mould. We are grateful to Michael Elowitz, Pau Formosa-Jordan, and Michael Kosicki for useful discussions. O.P. was supported by a Microsoft Research PhD Scholarship. Work in the Locke laboratory was supported by the European Research Council under the European Union’s Seventh Framework Program (FP/2007-2013)/ERC Grant Agreement 338060, a fellowship from the Gatsby Foundation (GAT3272/GLC), and an award from the Human Frontier Science Program (CDA00068/2012).

## Competing interests

The authors declare no competing financial interests.

